# Emotion induction modulates neural dynamics during ideational originality

**DOI:** 10.1101/2024.03.02.583080

**Authors:** Radwa Khalil, Sascha Frühholz, Ben Godde

**Author notes:** Corresponding author rsity. **Author Note**. Shared senior authorship (Sascha Frühholz, Ben Godde). We have no conflicts of interest to disclose.

## Abstract

Emotions remarkably impact our creative minds; nevertheless, a comprehensive mapping of their underlying neural mechanisms remains elusive. Therefore, we explored the influence of induced emotional states on ideational originality and its associated neural dynamics. Participants were randomly presented with three short videos with sad, neutral, and happy content. After each video, ideational originality was evaluated using the alternate uses task (AUT). Ideational originality was significantly higher after induction of the happy state than the neutral state; in contrast, there was a nonsignificant difference between the sad and neutral states. Associated neural dynamics were assessed through EEG time-frequency (TF) power and phase-amplitude coupling (PAC) analysis. Our findings suggest that emotional states elicit distinct TF and PAC profiles associated with ideational originality. Relative to baseline, gamma activity was enhanced after the neutral induction and more enhanced after the induction of a happy state but reduced after the induction of a sad state in 2-4 seconds after starting the task. Our PAC findings suggest that the attention system may be silent after the induction of a happy emotional state to load rich materials into working memory (WM) and active in the sad state to maintain these materials in WM.

**Highlight:**

1. Ideational originality was significantly higher after the induction of a happy state than in a neutral state.
2. Emotional states elicited distinct EEG time-frequency and phase-amplitude coupling profiles associated with ideational originality.
3. Relative to baseline, gamma activity was enhanced in the neutral state and more robust in a happy state but reduced in a sad state 2-4 seconds after starting AUT.
4. Enhancing ideational originality requires the induction of emotional states to suppress overlearned associations and strengthen weaker coupling associations, which is the case after the induction of a happy emotional state.

## Introduction

Creative thinking is a dynamic process that involves shifting between multiple modes of thought to generate novel and useful ideas ^1–6^. It is widely accepted that emotional states considerably influence creativity ^7–15^. However, whether positive or negative moods facilitate or inhibit creativity is an ongoing debate in the literature ^7–9^. While some studies support the perspective that positive affect improves various cognitive abilities, including creativity ^8,16^, other research suggests that a positive mood can inhibit creativity; on the contrary, a negative mood can facilitate it ^17,18^.

One of the most common operationalizations of creative cognition is divergent thinking (DT), which signifies a style of thinking that allows idea generation in a context where the selection criteria are relatively unclear, and more than one solution is correct; thus, DT implicates flexibility of mind ^19,20^.

Tests of DT are one of the most frequently used assessments of creative potential, and the alternate uses task (AUT)^21^ is among the most frequently used assessment methods. In the AUT, participants are requested to list alternative uses for everyday objects, such as bricks, clips, toothbrushes, newspapers, etc., and dimensions of DT, such as originality, fluency, flexibility, elaboration, etc., which could be calculated from the responses (for review, see ^22^). Several theories in creativity research have emphasized the significance of Originality in DT as a crucial component of creativity and effectiveness for an idea to be considered creative ^4,23–27^.

One commonly used operationalization approach to evaluating ideational originality is frequency-based (FB) scoring, which relies on the statistical infrequency of original ideas. The appeal of the FB technique lies in its objectivity when measuring originality ^26,28^, and this quantitative method has been used in numerous studies. For example, to examine the correlation between originality and dopamine (DA) activity, as indicated by the rate of spontaneous eye blinking (sEBR; ^23,29–32^, for review, see ^9^), and that DA is typically associated with a positive mood^7,8^.

There is growing evidence that the neural mechanisms of creative ideation do not depend on a single mental process or specific brain region as previously considered, i.e., asymmetric hemispheric activation ^33^, alpha synchronization ^34^, low arousal ^35^, or defocused attention ^36^. Nevertheless, providing an empirical integrative framework for the involved emotional and cognitive neural elements is still pending and should be systematically addressed ^9,37,38^. Regardless of how and where novel ideas are generated in the prefrontal cortex (PFC) and its brain circuits’ computations that convert them into creative behavior are performed by evaluating the novelty’s appropriateness and implementing its creative expression ^3,9,37,39–43^. Other neural circuits may be activated depending on the processing of information by the PFC circuits ^37,42^. These circuits could include sensory cortices (for visual and/or auditory information), emotional and associated limbic regions (for affective information), motivational systems (for appetitive or aversive information), and sensorimotor-related regions (for sensorimotor information) ^3,37,40,42,43^.

Brain dynamics and functional connectivity related to creative thinking are typically assessed with electroencephalography (EEG) ^38,44–47^ and functional magnetic resonance imaging (fMRI) ^6,40,48–53^. EEG allows a more fine-grained temporal analysis of brain activation in response to specific cognitive events, although it has considerably lower spatial resolution than fMRI ^45,47^. EEG studies on creative thinking have consistently revealed two principal neural features. The first implies increased task-related alpha-power in frontal brain areas during creative tasks compared with baseline ^44,46,54–57^. The second relates to a hemispheric asymmetry with a right-hemispheric dominance for alpha power ^33,47,55,58–63^. In line with these prior neurophysiological findings, it had been shown that bi-hemispheric activity in frontal sites and left-lateralized activity in central, temporal, and parietal sensor sites were predictors of an increase in originality ^44^.

Besides these two main features, researchers have identified other potential EEG biomarkers of creative thinking by adopting neural network analysis through cross-frequency coupling (CFC). CFC is a phenomenon where neural oscillations in distinct frequency bands interact within or across brain regions, giving rise to a complex regulatory structure ^64^. One type of CFC is phase-amplitude coupling (PAC), which describes the statistical dependence between the phase of a low-frequency brain rhythm and the amplitude or power of the high-frequency component of coupled electrical brain activity ^64,65^.

Previous studies support the executive nature of creativity ^10,41,66–70;^ for review, see ^22^. Accordingly, creative thought relies not solely on spontaneous thought processes but on controlled top-down activity ^49,71^. PAC effectively integrates activity across various spatial and temporal scales and plays crucial functional roles in neural information processing and cognition ^64,72^. The multifunctional process of creativity was revealed through PAC between theta, alpha, and gamma frequency bands ^73^. It has been proposed that CFC between parietal low-frequency (theta and alpha) and frontal high-frequency (gamma) bands is associated with subnetworks of working memory (WM), which allow for the maintenance and guidance of information over a short time ^74^, and this fronto-parietal CFC pattern might facilitate the generation of creative ideas ^75^. WM function is partly supported by frontoparietal activity as part of the executive control (EC) network ^76,77^.

The EC network is vital for directing attention to external stimuli ^78^, and it was suggested that functional connectivity between frontal and parietal cortices implies the involvement of attention in creative ideation ^39,49,57^. It was reported that pianists who expressed emotions increased FC between the dorsolateral PFC and the default network (DN) during improvisation^79^. The DN is notable for its activation in the resting state and is mainly involved in self-generated thought, which can be either spontaneous (mind-wandering) or goal-directed (mental navigation) ^42,49^. Interestingly, the two systems (EC and DN) are thought to have opposite effects on WM: frontoparietal activity is vital for directing attention to external stimuli^78^, while DN activity is crucial for internally directed cognition^80^. Therefore, nodes in the frontoparietal network (i.e., connector nodes), through diverse connections across brain systems and strong ties to each other, tune the connectivity between these two systems to achieve integrated cognition ^81–84^.

Our current study aims to comprehensively examine the influence of inducing emotional states on ideational originality and the associated neural network dynamics observed during ideational originality. After the positive, neutral, and negative emotional state induction, we evaluated original ideation with alternate uses task (AUT) ^21^. Participants were introduced to three short emotional videos in counter-balanced order, each eliciting a different emotion (happy, neutral, and sad). The ideational originality was then assessed with the AUT. The uniqueness of responses was assessed by employing the FB scoring method ^26,28^. During the AUT, EEG was recorded, and the associated neural network dynamics were evaluated using time-frequency (TF) power analysis and PAC analysis. We used these two analyses to determine how the frontal and parietal areas communicate over long distances during original ideation.

Based on the hedonic-tone hypothesis^8^ and the activating hypothesis ^85–87^, emotional states can either facilitate or hinder creative thinking because they are more activating than neutral conditions. Previous studies suggest positive emotions should boost creativity, while negative ones should hinder it ^7,13,86,88^. Therefore, we expected increased and decreased ideational originality in happy and sad emotional states, respectively, compared to the neutral state. Regarding the underlying neural mechanisms, our study was rather exploratory. We nevertheless expected that emotional states (positive, neutral, and negative) should exhibit distinct neural TF and fronto-parietal PAC signatures during original ideation, indicating how emotional states would affect information processing differently.

## Methods

### Participants, Ethical Approval and Eligibility Criteria

The study followed the Declaration of Helsinki (DoH) in all aspects. Participants were recruited from Constructor University, Germany, and received course credits. Before the study, participants completed a questionnaire about their mental and physical health, drug and medication use, and family history of disease before the experiment. Those with a history of diagnosed neurological disease or psychiatric disorders, heart conditions, severe head injuries, seizures (personally or in first-degree relatives), recurring syncopes, or learning disabilities were excluded.

All participants underwent eligibility screening for the EEG procedure and signed an informed consent form before starting the experiment. No participant had any previous experience with the AUT. The final sample consisted of 28 (14 male and 14 female) healthy undergraduate students between 18 and 23 years of age, and two participants were excluded from the EEG analysis. These two participants did not have clear EEG event markers in their recordings.

### Experimental Procedure

In three separate conditions, conducted in counter-balanced order, participants started with watching a 5-minute video with either sad, neutral, or happy emotional content to induce the respective emotions (cf. Figure 1). After each video, we provided the participants with the Self-Assessment Manikin (SAM) scale to rate their emotional arousal and pleasure levels and check the effectiveness of emotion induction. Then, we evaluated the creative ideation using AUT. We recorded the EEG during the AUT for time-frequency and subsequent functional connectivity analysis. In total, each condition lasted for about 15 minutes.

**Figure 1.**
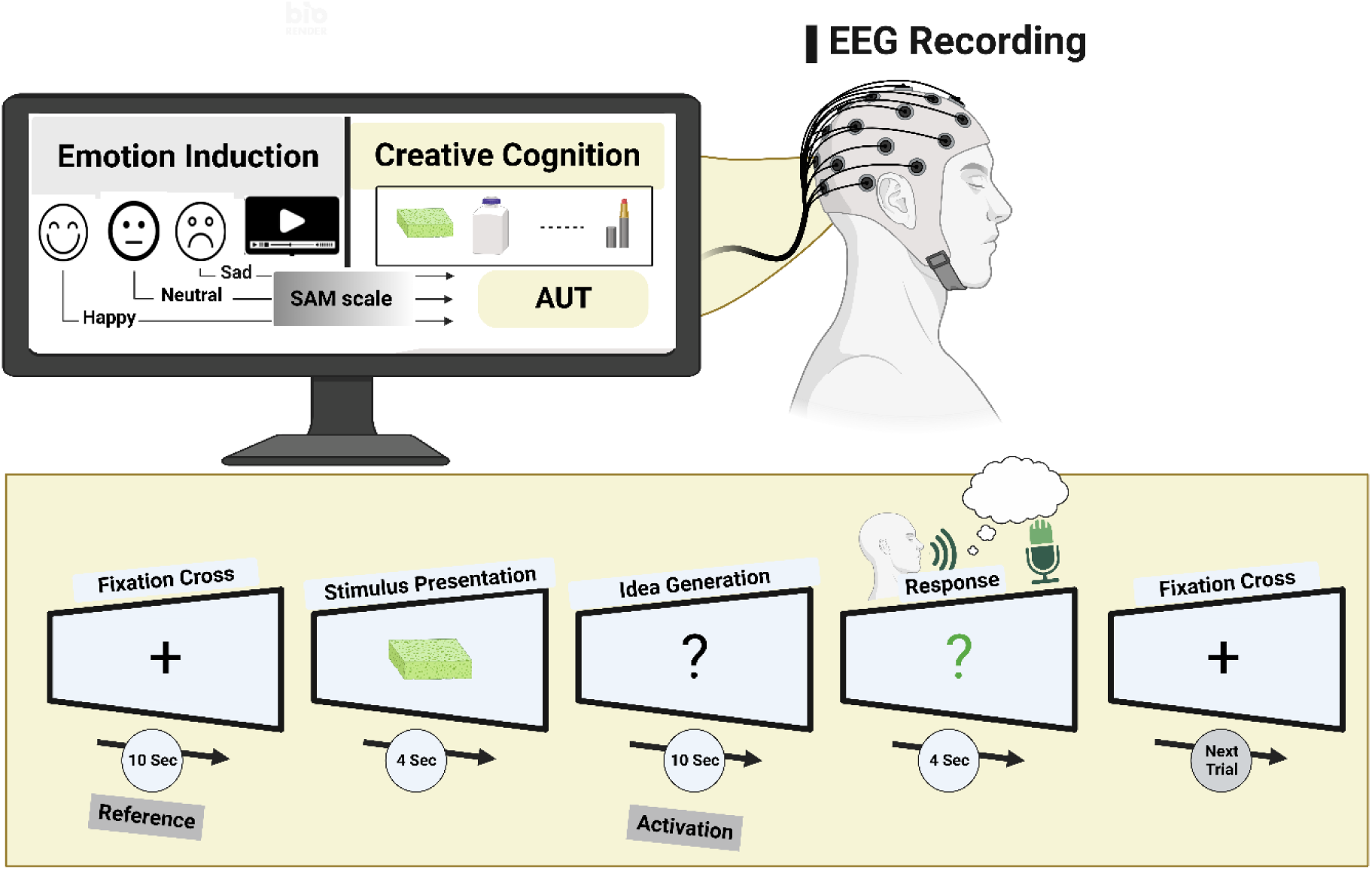
Illustration of the experimental setup. We introduced the participants to three short videos with different emotional contents—sad, neutral, and happy—for five minutes each in counter-balanced order. After each video, we provided the participants with the Self-Assessment Manikin (SAM) scale to rate their emotional arousal and pleasure levels and check the effectiveness of emotion induction. Then, we evaluated the creative ideation using the alternative uses task (AUT). The idea generation process was instructed to generate *original, untraditional uses* of stimulus presentation. The EEG was recorded during the AUT for time-frequency and functional connectivity analysis.

### Emotion Induction

Before starting the AUT, emotional states were induced through short films that lasted for five minutes each. These films were adapted from a past study ^14^ that induced positive, neutral, and negative emotional states. After each film, participants rated their current emotional states of arousal and pleasure using the 5-point SAM scale ^89^. The SAM is a well-validated, nonverbal pictorial assessment technique that directly measures the arousal, pleasure, and dominance associated with a person’s affective reaction to various stimuli.

The SAM scale was used to ensure the effectiveness of the emotion induction. The subjects rated their arousal and pleasure after watching each short film. The most positive effect was coded with the number 5, whereas the most negative effect was coded with the number 1. It ranged from a smiling, happy figure to a frowning, unhappy figure.

### AUT

In the AUT, everyday objects, for example, a sponge or bottle, were presented on the screen, and participants were asked to develop unconventional and original uses for these objects^10,41,70^; see appendix for examples of items used. Participants were carefully instructed on how to perform the AUT following the conventional task instructions^90,91^. The AUT started by presenting a fixation cross for 10 seconds (reference period; cf. ^46^).

Afterward, the stimulus picture of an everyday object appeared on the screen for 4 seconds. This stimulus was followed by a white question mark instructing participants to develop original ideas for using the given object over 10 seconds (the idea generation period). The idea generation process was instructed to generate *original uses* of stimulus presentation. Subsequently, the color of the question mark changed to green, signaling that the participants could articulate their ideas within 4 seconds. The oral responses were recorded and transcribed before the subsequent trial started. Fifteen items adapted from the experimental protocol by Schwab et al. (2014)^46^ were applied in each condition, i.e., after neutral, happy, and sad emotion induction. The presentation of the AUT stimuli was fully randomized, and we assured each participant that there was no repeated item to avoid any memory rumination of the items. We used the standard AUT scoring method from the Runco Creativity Assessment Battery (rCAB^4,28^

### Pre-Test

To create a valid criterion of response frequency, a group of 65 undergraduate students who did not participate in this study completed the AUT to predefine ideas’ originality, as Torrance (1974)^92^ suggested. They were given the same everyday objects used in this study. For each object, all original uses were collected using the same time constraints of the experiment. This list was used to compute a statistical infrequency measure, which was then used to assess the originality score for each response and, ultimately, for each participant. A score of zero was assigned to usage if it was mentioned by five percent or more of participants, one if it was mentioned by two to four percent of participants, and two if it was mentioned by less than one percent of participants. These statistical infrequency scores calculated an average originality score for each participant, ^93–95^.

Prior to calculating statistical infrequency, the database was appropriately modified to consider the similarity of responses based on the specified criteria: (i) extraneous words such as "used for" were removed; (ii) singular and plural words were made consistent; (iii) diminutives were avoided; and (iv) other minor adjustments were made to aid in identifying equivalent responses, for review see ^96^. Thus, an original response is statistically infrequent in the context of the selected sample.

### Statistical Analysis

We analyzed all data using Jamovi software ^97,98^. To test the effect of emotion induction on arousal and pleasure, we conducted one-way ANOVAs with the mode of emotion induction (neutral, happy, and sad) as the within-subjects factor and arousal and pleasure ratings as dependent variables, followed by post-hoc comparison tests. Similar one-way ANOVA was performed with the Z transform of the ideational originality as the dependent variable, followed by a post-hoc comparison test to test the effect of emotion induction on originality. Results were interpreted as statistically significant if the *p*-value was <.05, corrected for multiple testing where applicable. For data visualization in Figures 2 and 3, we used Graph Prism (GraphPad Software, San Diego, California USA, www.graphpad.com).

**Figure 2.**
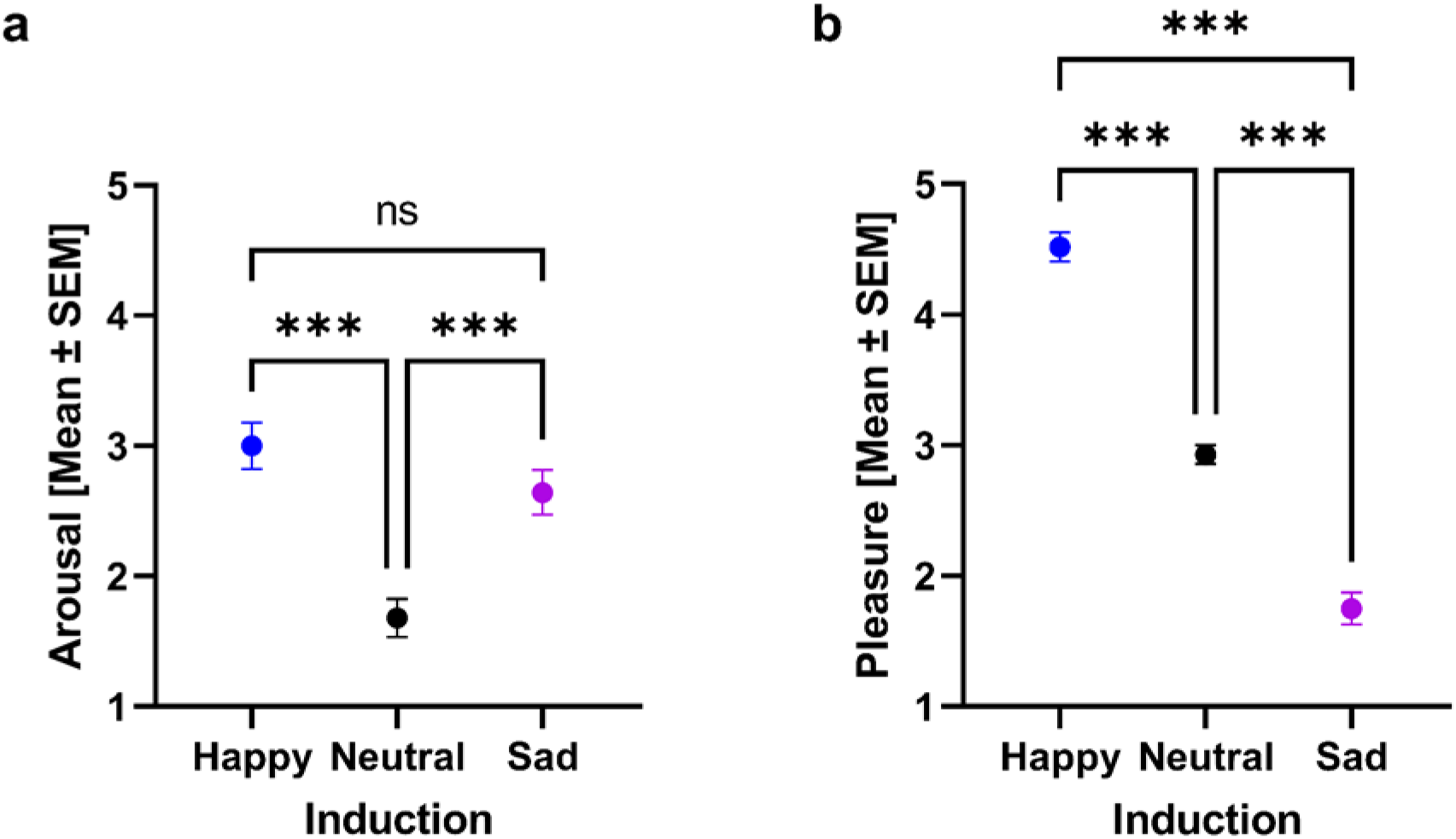
Effectiveness of emotion induction elicited through short films with neutral, happy, and sad emotional content. Panels a and b refer to the effects on arousal and pleasure. The y-axes of each panel refer to the mean and standard error of the mean [SEM] for arousal and pleasure after each emotion induction as represented on the x-axes; for detailed statistics, see Tables 1 and 2. Stars indicate significance (\**p* < .05. \*\**p* < .01. \*\*\**p* < .001).

**Figure 3.**
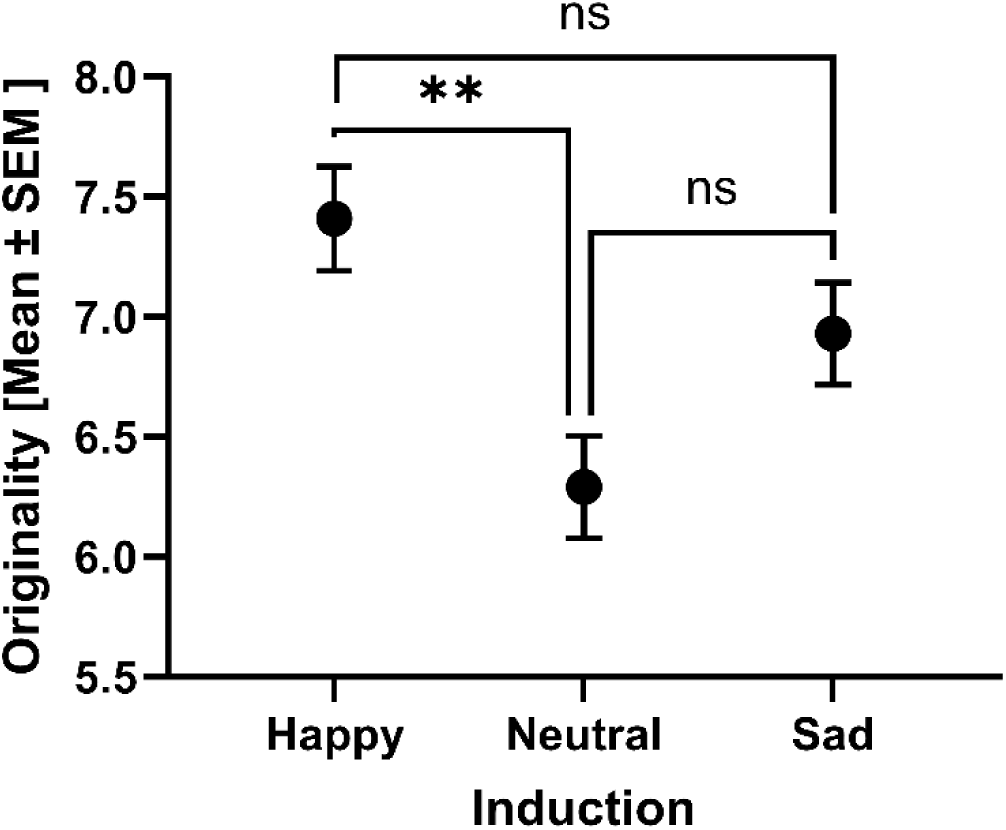
Effect of emotion induction on originality index. Emotion induction (happy, neutral, and sad) affects originality. See Table 3 for detailed statistics. Stars indicate significance (\**p* < .05. \*\**p* < .01. \*\*\**p* < .001).

**Table 1.**
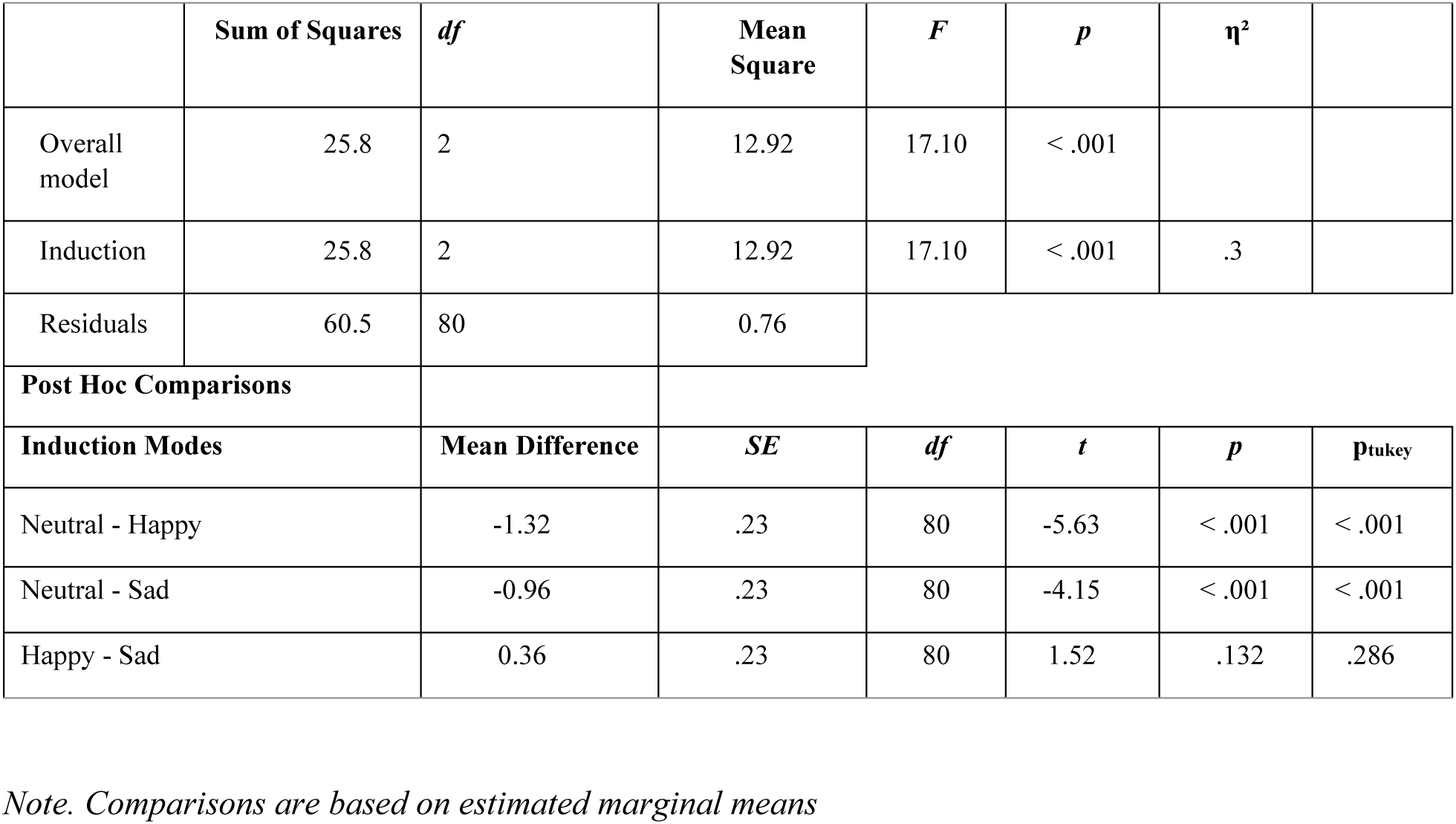
ANOVA of emotion induction elicited through short films using SAM scales for arousal.

**Table 2.**
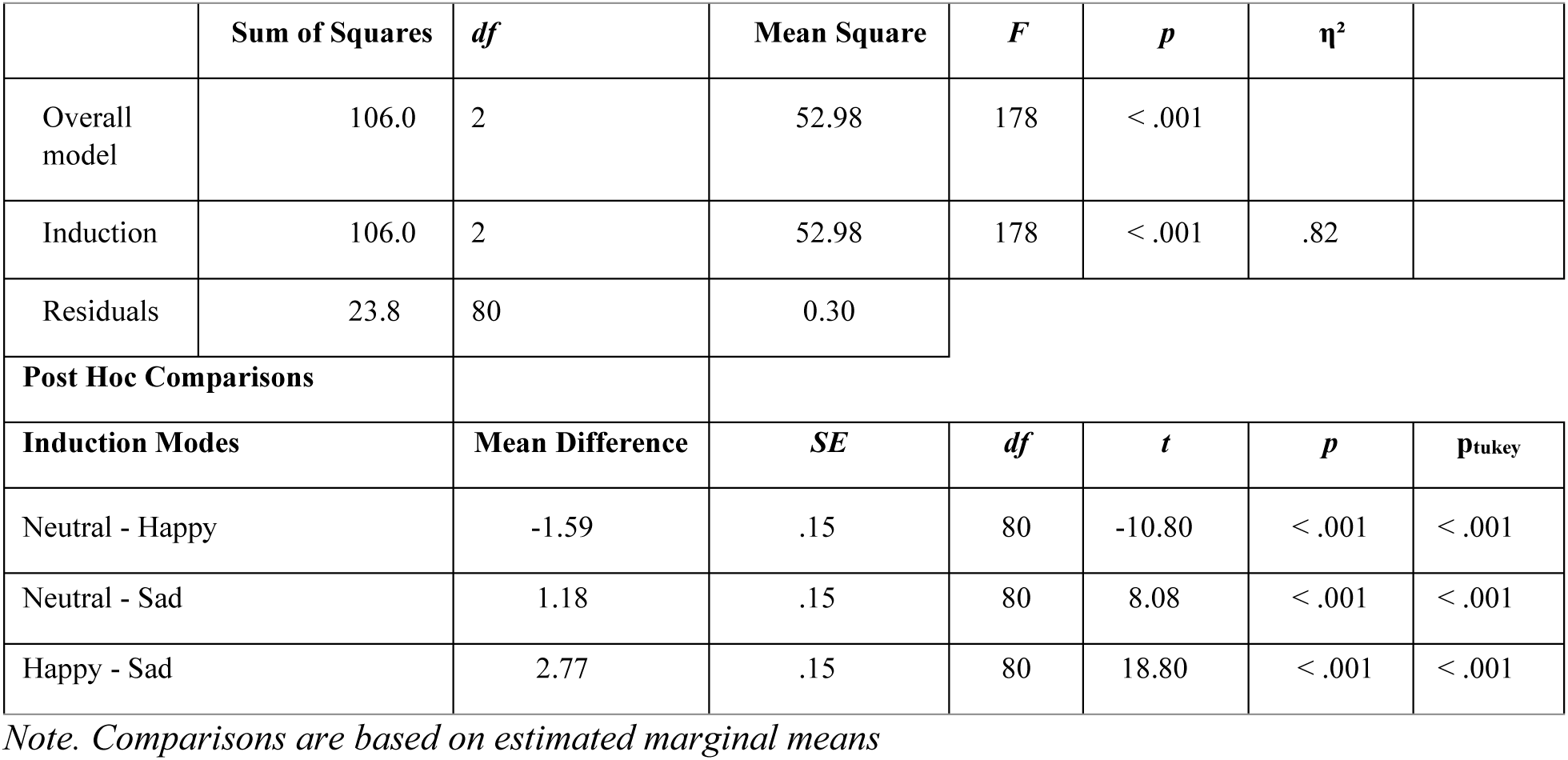
ANOVA of emotion induction elicited through short films using SAM scales for pleasure.

**Table 3.**
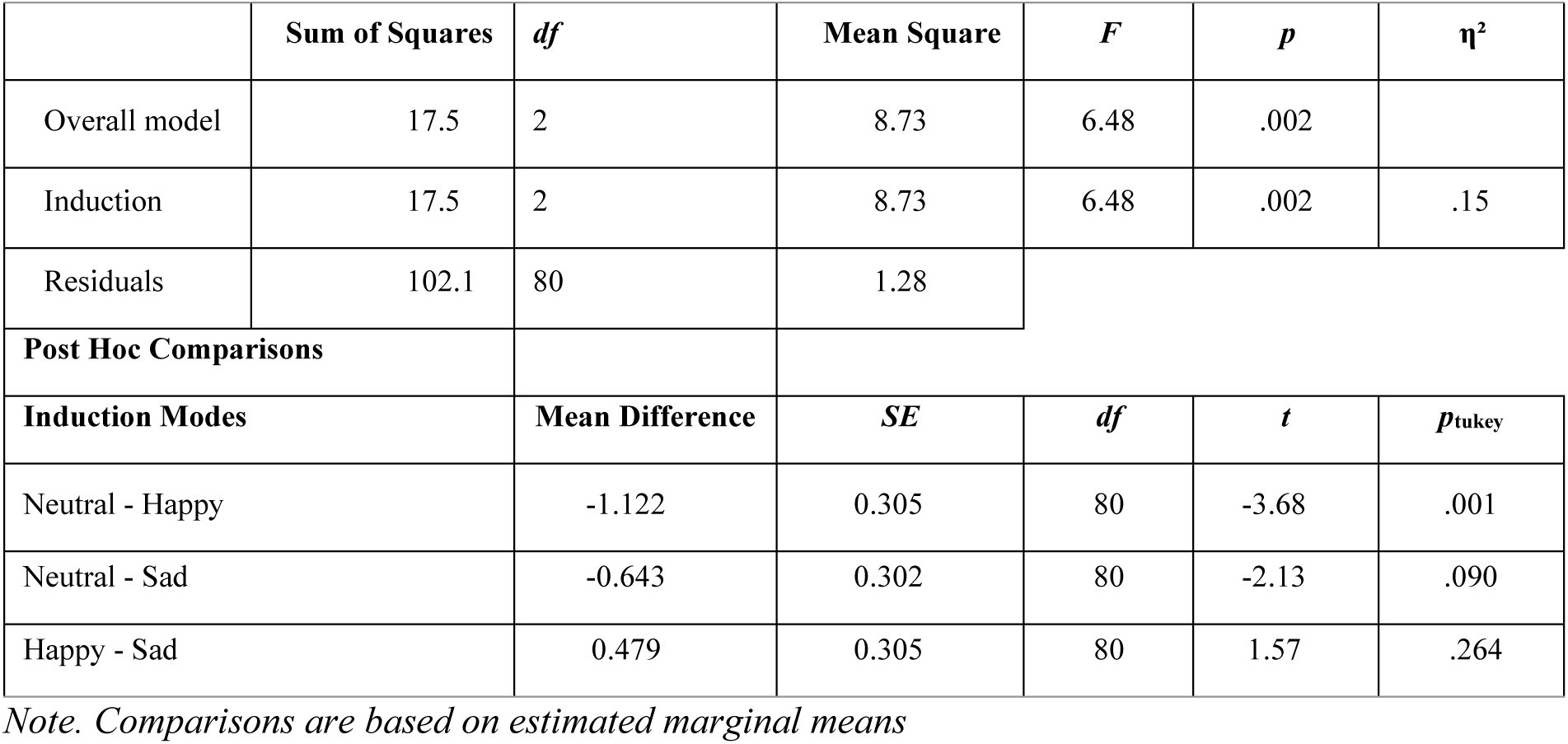
ANOVA of the effect of emotion induction on originality.

| | Sum of Squares | df | Mean Square | F | p | $\eta^2$ |
| --- | --- | --- | --- | --- | --- | --- |
| Overall model | 17.5 | 2 | 8.73 | 6.48 | .002 |  |
| Induction | 17.5 | 2 | 8.73 | 6.48 | .002 | .15 |
| Residuals | 102.1 | 80 | 1.28 |  |  |  |
| **Post Hoc Comparisons** |  |  |  |  |  |  |
| Induction Modes | Mean Difference | | SE | df | t | $p_{tukey}$ |
| Neutral - Happy | -1.122 |  | 0.305 | 80 | -3.68 | .001 |
| Neutral - Sad | -0.643 |  | 0.302 | 80 | -2.13 | .090 |
| Happy - Sad | 0.479 |  | 0.305 | 80 | 1.57 | .264 |

*Note. Comparisons are based on estimated marginal means*

### EEG Recording and Offline Processing

EEG was measured simultaneously with the AUT. A 32-channel electroencephalogram (EEG) was recorded using Ag-AgCl electrodes mounted on an elastic cap according to the 10–20 electrode system, with a reference electrode at the nose tip (Nz). The EEG signal was amplified by a REFA multi-channel system (TMS International; www.tmsi.com) and digitized at a sampling rate of 512 Hz. Impedances were kept below 10 kΩ. A horizontal and vertical electrooculogram (EOG) was recorded at the outer canthi of the eyes, and both supra- and suborbital positions of the right eye to monitor eye blinks and movements.

We used EEGLAB software (version 2019.1; https://sccn.ucsd.edu/eeglab/index.php) for offline preprocessing and time-frequency decomposition of the EEG data. The EEG signal was then re-referenced to an average across the 32 EEG electrodes and down-sampled to a 256 Hz sampling rate during offline preprocessing. The signals from channels with large signal artefacts and signal drifts were reconstructed using the spherical interpolation method from neighboring electrodes (one electrode for six participants and two electrodes for one participant). We applied filtering to the EEG signal by first applying a notch filter to account for the 50 Hz direct current effects (47–53 Hz). We afterward applied a bandpass filter in the 0.1–100 Hz range. Blink-artifact detection and correction were performed using a spatially independent component analysis (ICA; infomax algorithm) implemented in the EEGLAB software. The EEG data were epoched around references and activation period onsets in a broad prestimulus and poststimulus time range (-8 to +18 s).

Time-frequency (TF) EEG analysis was applied to identify relevant frequency bands and their temporal changes during encoding. For this purpose, a continuous wavelet transformation was performed on single epoched trials for each subject, and four different clusters of electrodes over the left frontal (F3, FC5, FC1), right frontal (F4, FC6, FC2), left parietal (P3, CP5, CP1), and right parietal (P4, CP6, CP2) areas. The TF procedure allows analyses of all oscillatory activity, phase-locked and non-phase-locked, to the stimulus onset. The neutral state was characterized by increased gamma between 2 and 4 seconds after trial onset relative to the reference period, particularly in the left hemisphere, followed by gamma decrease in both hemispheres. The signal from each electrode and epoch was convolved with complex Morlet wavelets as follows: w(t, f0) = A exp(−t2/[2σ2t]) exp(2iπf0t), with σf = 1/[2πσt]. The wavelets were normalized such that their total energy was 1. The central frequency f0 ranged between 1 and 70 Hz, with the ratio f0/σf = 7. Subsequently, the time-frequency energy distribution of the reconstructed signal was obtained using Morlet wavelet expansion. The log-scaled values obtained for each trial (each electrode cluster and each frequency) were normalized to the mean baseline energy level from –3000 to 0 ms before trial onset, revealing the percentage decrease or increase of the in-band power during the encoding phase. The mean TF signal for the reference and activation periods was calculated separately for each electrode cluster and emotion condition for each participant.

Next, the participants’ mean TF signals were averaged across participants for each electrode cluster and emotion condition. The statistical significance of the relevant time and frequency areas was determined by a permutation test (2,000 permutations), specifically by randomly shuffling time points and frequency bands for participants’ mean TF signals within an electrode cluster, experimental period, and emotion condition. For all permutations, we calculated the difference in TF signals between the activation and the reference period and smoothed the TF signal by averaging over neighboring TF matrix elements (the center TF element and the first and second neighboring TF elements) to obtain a distribution of 2,000 values for each TF signal of the four electrode clusters and three emotion conditions. The grand average and smoothed TF difference between the activation and the reference period were compared against this distribution of 2,000 values using a normal cumulative distribution function. A time-by-frequency area was determined to be significant using a two-sided test and a p <.001. An additional cluster threshold was applied such that only TF areas with k >= 100 neighboring significant TF areas were determined to be significant.

We performed a CFC analysis between the two frontal electrode clusters and the two parietal electrode clusters. This CFC analysis was set up to determine the coupling between the 1–12 Hz range (delta: 1–3 Hz, θ: 3–7 Hz, alpha: 7–12 Hz) as the frequency phase range that convolved the amplitudes of the higher frequency in the low (30–50 Hz) and high (50–70 Hz) ranges. We computed the CFC analysis using the Tensorpac toolbox (version 0.6.5; https://etiennecmb.github.io/tensorpac/) and the mean vector length method^99^, permuting the phase across trials^100^, and normalizing data by subtracting and dividing by means of the surrogates. All CFC analyses were performed separately for each participant, and the resulting CFC data was smoothed the same way as the TF data before we averaged them across participants. We used the same permutation procedure for the TF data to determine CFC areas of significance. A two-sided test determined significance with p <.001 and k ≥ 12 for a significant number of neighboring areas.

### Ethical Approval

The authors state that this study was approved by the local ethics committee of Constructor University and has followed the principles outlined in the Declaration of Helsinki for human experimental investigations. Informed consent has been obtained from all the participants.

## Results

### Emotion Induction

To check the effectiveness of the emotion induction, participants rated their emotional arousal and pleasure levels after each video with the Self-Assessment Manikin (SAM) scale ^89^. An analysis of variance using general linear modeling revealed that emotion induction through videos with emotional content was successful.

ANOVA with the mode of emotion induction (neutral, happy, and sad) as the within-subjects factor and *arousal* as the dependent variable revealed a significant main effect of induction mode (*F (2,80) = 17.1*; *p <* .001, *p η² =*.30; cf. Figure 2a and Table 1 for detailed statistics).

The highest level of arousal was observed in the happy induction mode, while the lowest was in the neutral induction mode. Post-hoc analysis confirmed significant differences between neutral and happy emotion induction modes (*t =* –5.63, *p_Tukey_ <* .001, df = 80) and neutral and sad emotion induction modes (*t = –*4.15, *p_Tukey_ <* .001, df = 80). However, there were no significant differences between the happy and sad emotion induction modes (*t =* 1.52, *p_Tukey_=*.29, df = 80).

Similarly, ANOVA with a mode of emotion induction as the within-subjects factor and *pleasure* as the dependent variable revealed a significant main effect of induction mode (*F* (2,80) = 178, *p <* .001, *η²* =.82; cf. Figure 2b and Table 2 for detailed statistics). The highest level of pleasure was for the happy induction mode, while the lowest was for the sad induction mode. Post-hoc analysis confirmed a significant difference between neutral and happy induction modes (*t =* –10.80, *p_Tukey_ <* .001, df = 80), neutral and sad induction modes (*t =* 8.08, *p_Tukey_ <* .001, df = 80), and happy and sad induction modes (*t =* 18.80, *p_Tukey_ <* .001, df = 80).

### Effect of Emotion Induction on Ideational Originality

Figure 3 illustrates the effects of emotion induction on originality. ANOVA was performed with emotion induction mode as the within-subjects factor. Originality scores were used as dependent variables (cf. Figure 3 and Table 3 for detailed statistics).

ANOVA revealed a significant effect of emotional induction mode (*F (2,80) = 6.48; p =.002, η² =.15*). In contrast to the neutral condition, originality was enhanced in the happy condition but not the sad one. Post-hoc tests confirmed significant differences between the neutral and happy induction modes *(t = -3.68, p_Tukey_* =.001, df = 80), but no significant differences were recorded between the neutral and sad *(t = -2.13, p_Tukey_* =.090, df = 80) or between the happy and sad induction modes *(t = 1.57, p_Tukey_* =.264, df = 80).

### EEG Time-Frequency (TF) Analysis and Cross-Frequency Coupling (CFC) During AUT

We conducted TF analysis with continuous wavelet transformation on single-epoched trials separately for reference and activation periods for each subject and four different clusters of electrodes. These clusters spanned over the left frontal (LF: F3, FC5, FC1), right frontal (RF: F4, FC6, FC2), left parietal (LP: P3, CP5, CP1), and right parietal (RP: P4, CP6, CP2) areas (Figure 4a).

**Figure 4.**
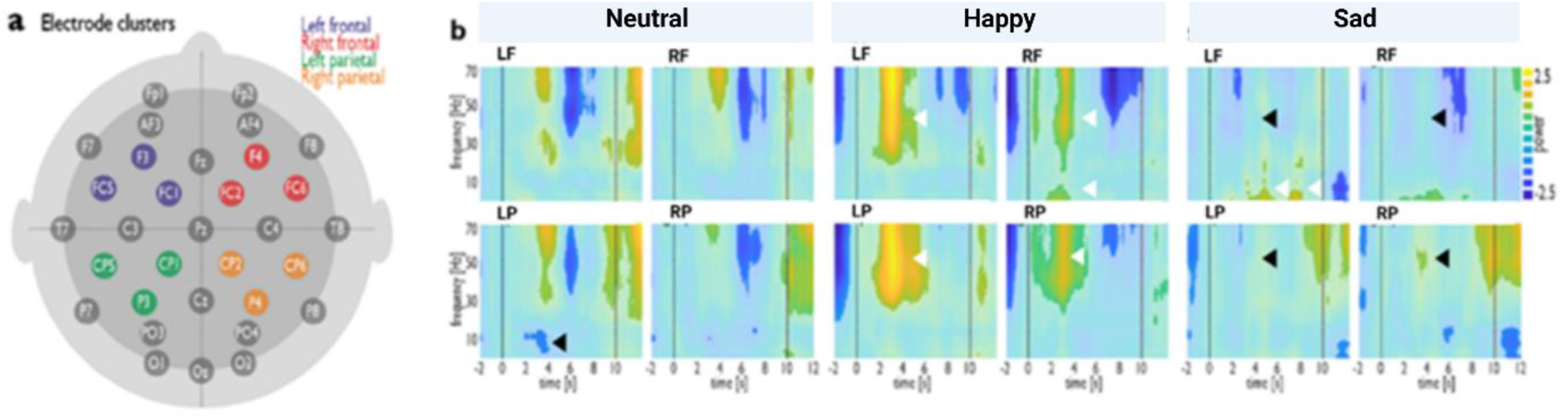
Time-frequency (TF) plots for EEG data obtained during AUT trials after each emotional induction (neutral, happy, and sad) Panel (a) refers to TF analysis with continuous wavelet transformation, which was performed on single epoched trials for each subject and four different clusters of electrodes over the left frontal (F3, FC5, FC1), right frontal (F4, FC6, FC2), left parietal (P3, CP5, CP1), and right parietal (P4, CP6, CP2) areas. Panel (b) illustrates TF plots for each emotion induction condition represented in the left frontal (LF) and right frontal (RF) areas (upper row) and left parietal (LP) and right parietal (RP) areas (lower row). Each TF plot displays the frequency in Hz (the power fluctuates from 2.5 to -2.5 as signified by the color scale) and the time in seconds on the y and x axes, respectively. Time 0 to 10 seconds refers to the AUT trial window, marked by vertical lines, and the prominent differences between conditions are indicated using arrows.

Each emotional induction revealed a distinct TF profile during AUT (Figure 4b). Reduced low-frequency (delta, theta, and alpha) power was observed in the LP cortex (Figure 4b). The happy state was characterized by a stronger gamma power increase relative to baseline in all four cortical regions but most prominent in the left hemisphere, followed by a less strong gamma reduction, particularly in the LP cortex. There was also an increase in low-frequency (delta, theta) power in the RF cortex 2-4 seconds after trial onset (Figure 4b). Conversely, there was nearly no change in gamma power in the sad state from the reference to the task period in all four cortical regions; instead, increased low frequency (theta, delta, and alpha) power could be observed in the LF cortex (Figure 4b).

The PAC patterns indicating the cross-frequency coupling between frontal and parietal regions of both hemispheres (as hubs of the EC networks in creative ideation) during DT after emotion induction are illustrated in Figure 5.

**Figure 5.**
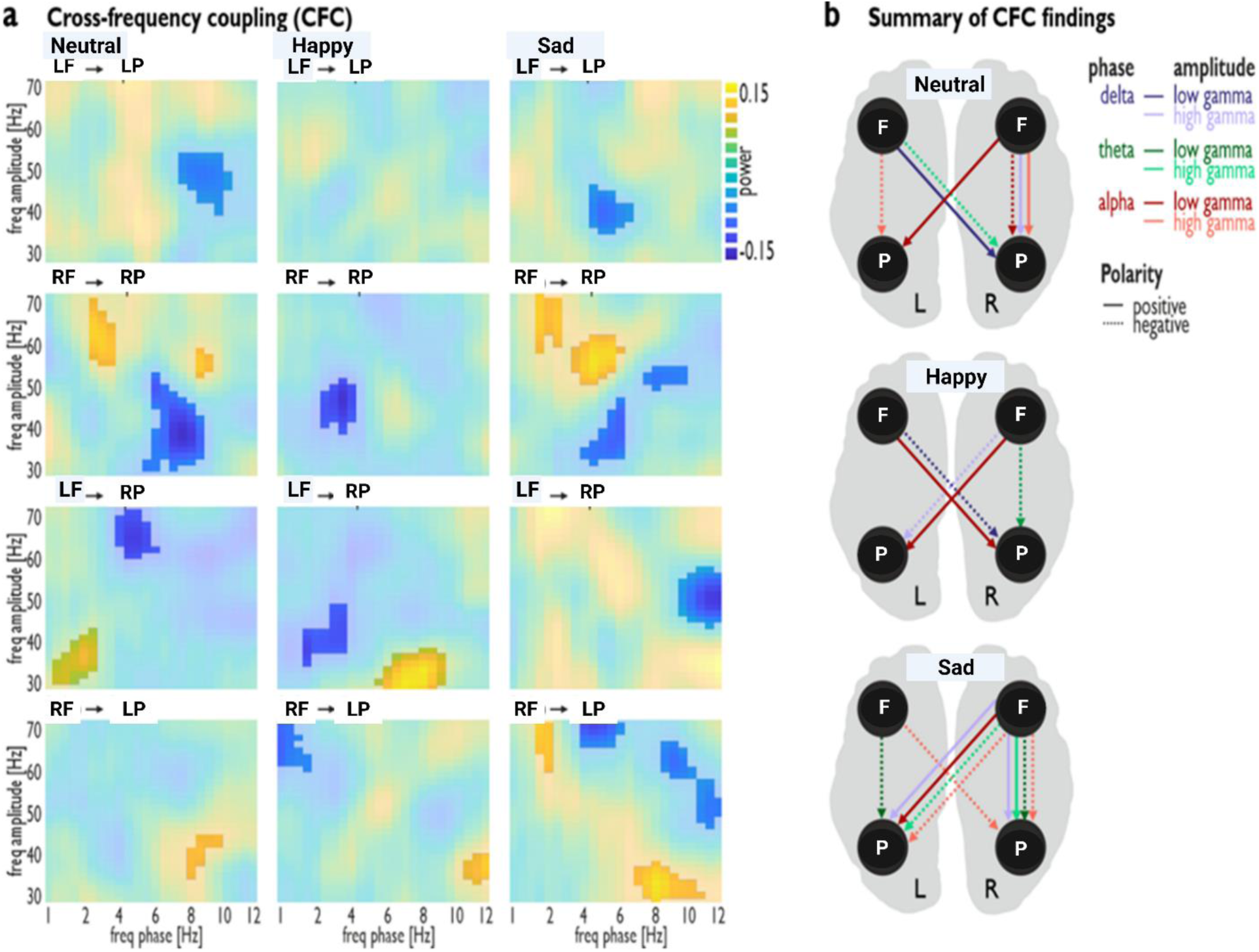
Cross-frequency coupling (CFC) during AUT after three emotion induction conditions (neutral, happy, and sad). Phase-amplitude coupling (PAC) paths (rows) are shown for each emotion induction condition (columns) as follows: left frontal to left parietal (LF–LP), right frontal to the right parietal (RF–RP), left frontal to the right parietal (LF–RP), and right frontal to left parietal (RF–LP) cortex. Each spectrogram displays the frequency of the phase signal in LF and RF on the x-axes and the frequency of the amplitude signal in LP and RP on the y-axes with its normalized PAC value (range: +0.15 to -0.15; cf., color scale). The PAC value is a normalized measure of how much the phase of the lower-frequency component modulates the amplitude of the higher-frequency component. Positive values indicate enhanced modulation, while negative values indicate reduced modulation.

PAC values represent the coupling strength between the phase of the low-frequency oscillations (delta: 1–3 Hz, theta: 3–7 Hz, alpha: 7–12 Hz) in the frontal cortex and the amplitude of parietal lower (30–50 Hz) and higher (50–70 Hz) gamma oscillations relative to all other factors that modulate the high-frequency amplitude. Positive PAC values indicate a more robust modulation of parietal gamma by the phase of frontal, low-frequency oscillations (i.e., stronger coupling) during the task as compared to baseline. On the contrary, negative PAC values indicate weaker modulation of parietal gamma oscillations by the phase of frontal, low-frequency oscillations (i.e., weaker coupling).

Remarkably, a contralateral path with positive polarity, which is right frontal (alpha)-left parietal (low gamma), was maintained for the three conditions. However, each induced condition has its distinct profile, as illustrated in the summary of CFC findings in Fig. 5, which describes the coupling pathways in each induced condition.

Relative to the reference period, the neutral condition was characterized by three contralateral couplings and four ipsilateral couplings with different polarities, i.e., positive and negative. The two positive contralateral couplings indicate frontal (alpha and delta) - parietal (low gamma) paths. The negative contralateral coupling directs the frontal (theta)-parietal (high gamma) path. The two positive ipsilateral paths refer to frontal (delta)-parietal (high-gamma) and frontal (alpha)-parietal (high-gamma). The two negative ipsilateral relate to frontal (alpha)-parietal (low and high gamma) paths.

The happy condition was characterized by four contralateral couplings with different polarities and one ipsilateral coupling with negative polarity. The two positive contralateral couplings indicate frontal (alpha) - parietal (low gamma) paths. The two negative contralateral couplings refer to frontal (delta) -parietal (low and high gamma) paths. The negative ipsilateral path couples the frontal (theta)-parietal (low gamma). The sad condition was characterized by five contralateral and five ipsilateral couplings with different polarities. The two positive contralateral couplings show frontal (alpha and delta) - parietal (low and high gamma) paths. The three negative contralateral couplings refer to frontal (alpha and theta)-parietal (high gamma) paths. The two positive ipsilaterals relate to the frontal (delta and theta)-parietal (high gamma) path. The three negative ipsilaterals relate to the frontal (theta and alpha) -parietal (low and high gamma) paths.

## Discussion

Our findings shed light on the impact of affective states (neutral, sad, and happy) on ideational originality and, for the first time, provide the related neural dynamics as revealed by TF and PAC profiles. We summarized the effects of emotion induction on creative ideation and PAC in Figure 6.

**Figure 6.**
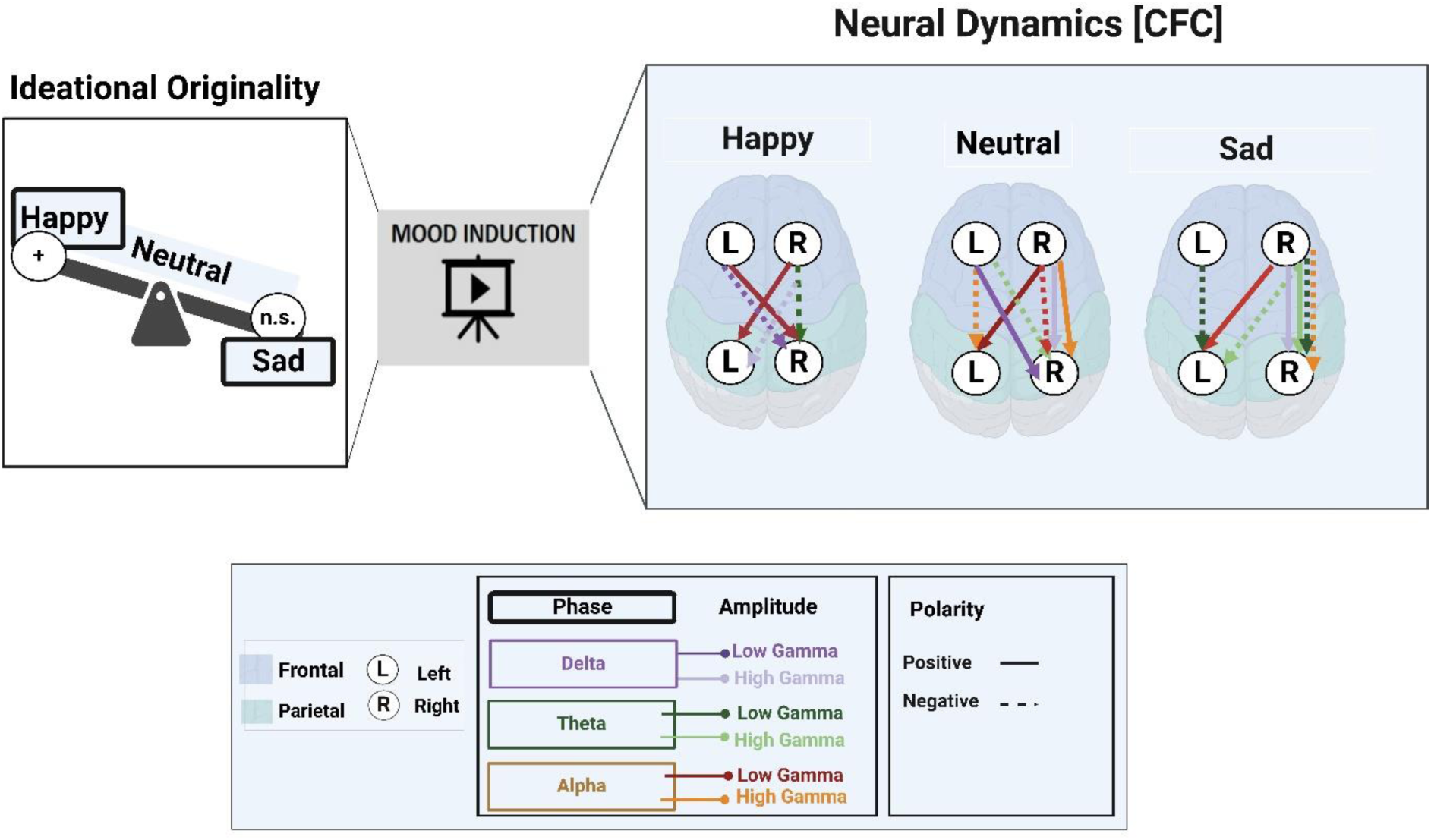
Summary of effects of emotion induction on both the original ideation and Cross-Frequency Coupling (CFC). The behavioral level represents the selected creative ideation (i.e., original ideation) that expresses a significant difference between happy and neutral states—the neural dynamics as a function of PAC in the happy, neutral, and sad states. The indications of the phase, amplitude, and coupling are described in the legend.

At the behavioral level, subjective ratings of arousal and pleasure revealed the effective induction of emotional states and their influence on ideational originality, which aligns with the activating hypothesis ^85–87^. Our behavioral result indicated that ideational originality was significantly higher after the induction of the happy state compared to the neutral state; in contrast, it was nonsignificant after the induction of the sad state compared to the neutral state. This behavioral finding aligns with the asymmetrical effects of emotional induction on ideational originality and supports the hedonic-tone hypothesis ^8^; positive emotional states facilitated ideational originality, while negative emotional states constrained it^7,13,86,88^. Enhancing the level of ideational originality after happy state induction might imply an individual’s fun or joy in a positive mood compared to a negative mood state^101^. In the same vein, previous research reported a relationship between DA and ideational originality ^23,29–32^ and that DA is typically associated with a positive mood, facilitating creative ideation ^7,8^.

### Neural Dynamics

Concerning neural dynamics, emotional states elicited independent and distinctive neural TF and fronto-parietal PAC profiles during the original ideation process. These distinct profiles reflect the asymmetric effects of the positive, neutral, and negative states on underlying information processing of ideational originality.

## 1. TF profiles

The happy state was characterized by a considerably stronger gamma increase in the early time window, 2–4 seconds after trial onset. This enhanced gamma activity might suggest the successful execution of WM^102^, and its transient nature with the subsequent decrease might reflect the prompt engagement in solving insight problems^103^. Despite the general dominance of the gamma signature in the left and right frontal and parietal cortices (LF, RF, LP, and RP) after the induction of the happy state, it was more robust in the left hemisphere, suggesting a bias toward the left hemisphere over the right hemisphere for WM loading during original ideation. This higher WM load might align with a dominance of alpha activity in the left hemisphere^60^. However, an increase in low-frequency oscillations (theta and delta) was observed only in RF. The theta-band activity might reflect the demand of RF for top-down control ^99^, and the delta-band activity might coordinate large-scale networks^104^.

In the sad emotion state, there was nearly no gamma activity in all four cortices (LF, LP, RF, and RP). The emergence of low-frequency oscillations in the frontal cortices (theta, delta, and alpha in the LF) could suggest the need for top-down control, directed attention, and multiple sensory modalities during WM maintenance ^74^. Notably, the increase in alpha activity might result from an increase in the magnitude of inhibitory bouts, which break down ongoing gamma activity ^105^for gating sensory inputs ^106,107^.

## 2. PAC profiles

A common coupling path was preserved in all three emotional states: the coupling of low gamma activity in LP to the phase of the contralateral RF alpha. Traditionally, PACs between the alpha phase and low gamma amplitude are called "sharp waveforms" ^108^. The maintenance of these sharp waveforms in all three emotional states during ideational originality supports the notion that alpha–gamma PACs are necessary for WM function^74^, gating sensory information ^105^, imagined actions^109^, and blocking out distraction^110^. Given the proposed role of the frontal regions in monitoring WM storage in distant sensory areas ^111,112^, it could be speculated that this frontal region controls high-frequency gamma activity in the sensory area of the parietal cortex.

These distinctive PACs might be interpreted as happy emotion induction being comparatively biased toward the spontaneous (automatic) processing mode instead of the deliberate processing mode during ideational originality. The execution of the automatic process requires unconscious thoughts that are relatively random, unfiltered, and unusual to be loaded in WM while silencing the attentional system in deciding for the content to become conscious ^37^. Moreover, alpha-gamma (i.e., "sharp waveforms") versus theta-gamma (i.e., "nested oscillations") codes for different WM information than theta-gamma (nested oscillations) ^74^. Theta-gamma coupling (also called "nested oscillations"^108^) has been suggested to be crucial for fostering WM and long-term memory by facilitating neural communication and supporting neural plasticity^113^, while delta-high-gamma PAC is implicated in attention^74^. Furthermore, an association of increased PAC between the frontal delta and theta and parietal gamma with WM maintenance has been suggested ^114,115^.

The lower number of PAC paths for the happy state might be interpreted as there is accessibility to a more extensive, diverse range of information and probably need fewer fronto-parietal couplings for EC, and that happy state might activate more material from WM ^116^. This might also explain a tendency to form fewer and simpler neural coupling paths reflecting better associations between ideas and differentiate between items and content more rapidly, resulting in a significantly higher level of ideational originality on the behavioral level than neutral induction. Conversely, the induction of a sad state represents less effective retrieval cues for material in WM, which may limit ideational originality and necessitate stronger and more likely coupling paths. These distinctive PAC patterns after sad induction might be interpreted as the sad state being comparatively biased toward the deliberate (control) processing mode instead of the automatic processing mode during the original ideation.

## Summary

Our findings suggest that the induction of happy and sad emotional states favor two facets of WM associated with varying modes of processing—deliberate and spontaneous—for ideational originality^37^. Consequently, the attention system might be silent after the induction of a happy state to allow the loading of rich materials into WM; in contrast, it becomes active after the induction of a sad state for maintaining items in WM ^116^, and this is reflected in the association of increased PAC between the frontal delta and theta and parietal gamma with WM maintenance, as has been suggested ^114,115^. The emergence of higher-level ideational originality after emotional induction of a happy state might suppress overlearned associations and enhance weaker coupling^108^, while delta-high-gamma PAC is implicated in attention ^74^.

To the best of our knowledge, this study is the first to report the influence of three basic emotional inductions on the creative ideation process of DT as a function of ideational originality and associated neural dynamics. However, our current investigation will be part of the foundation for further neurocognitive, behavioral, and brain imaging research on emotional induction and DT to be extended in future studies. Therefore, we highlighted the limitations of this study and the future directions in the coming section.

### Limitations, Open Questions, and Future Directions

The past decade has witnessed a rise in interest in the impact of emotional affective states on creative ideation; however, these neuroscientific investigations and related neural dynamics remain open for further investigation. The interpretations and discussions of our findings should be read in light of their limitations and noted as such. This circumstance was unavoidable because no prior study explored the influence of induced emotional states on ideational originality and its associated neural dynamics using TF and PAC.

Here are examples of the limitations of our current study. Due to the within-subject design, there might still be effects on the order of the videos. Therefore, ruminations about the sad condition are possible during the neutral condition. Although we used the SAM scale after each video to measure arousal and pleasure levels, indicating the induction’s effectiveness, we could not rule out the possibility of the participants getting bored or having mixed feelings. Furthermore, we have a relatively small sample size of university students, which should be replicated in the future with a larger sample size.

The other point is that the imposed time constraint during EEG in our experiment may have limited the behavioral scoring of ideational originality and did not allow for measuring fluency and flexibility. For instance, participants had to generate original alternative uses for the presented objects in a limited time. However, originality requires time to emerge, so only low levels of originality can be captured in this short experiment duration (i.e., 10 seconds). Also, the task was instructed to generate original ideas (i.e., original uses rather than alternative uses, which could allow for more ideas that could be counted for fluency and flexibility). Therefore, expanding our findings by replicating this experiment using longer response durations and allowing participants to provide more original responses is beneficial. Likewise, the instruction could be changed to alternative uses so that it is not only restricted to uniqueness or original ideation but also to other dimensions such as fluency and flexibility.

In this study, we statistically analyzed EEG data only at the group level, and no individual data that could be associated with individual behavioral data were extracted. However, addressing this issue in light of individual differences would provide another perspective for future research.

Although the FB originality method used in this study is a valid scoring approach, determining the frequency of responses categorized as "uncommon" requires subjective assessments and adjustments to determine if specific responses are similar or distinct from one another ^96,117^. Given that Originality in DT tasks encompasses dimensions of uncommonness, remoteness, and cleverness ^118,119^ thus, it is beneficial to expand this study in the future by incorporating rater-based scores to capture the dimensions of remoteness and cleverness in addition to uncommonness.

In this study, we focused only on the induction of happy and sad emotional states, and it is worth extending this study to focus on other specific feelings, for example, joy, anger, and contentment, and to provide a broader framework for the relationship between emotion induction and ideational originality and its underlying neural dynamics. A crucial facet to consider in future research is the intensity of emotion, as previous research has shown that optimism intensifies positive emotions (i.e., enthusiasm and happiness) and diminishes negative emotions (i.e., sadness and fear) ^7,88^. In the same vein, extending this study by exploring the effect of sEBR changes as reflections of phasic DA changes on ideational originality in association with the underlying dynamical neural mechanisms would be noteworthy.

When interpreting our study’s results, it is worth considering that we focused on young adults’ everyday creativity; thus, our results are likely more typical for young people than for more professionalized creative functioning^120^, and our results differ when compared with figural measures of creativity^121^. According to a recent meta-analysis^121^, several studies have also found that the diverse types of DT tasks assess slightly different aspects of DT, especially when comparing verbal and figural measures^122–126^ .

Future studies should include both EC-related task(s) and creative ideation tasks (i.e., AUT) to strengthen the generalizability of our current findings. It is also valuable to consider intelligence components, as previous research has indicated strong connections between DT and intelligence components ^66,127–130^.

## Author Contributions

RK, SF, and BG designed the research and the experimental design. RK conducted the experiment, RK and BG performed the behavioral analysis, and SF performed the time-frequency and functional connectivity analysis. RK drafted the manuscript with SF and BG’s significant insights.

## Funding Acknowledgements

SF received support from the Swiss National Science Foundation (SNSF PP00P1_157409/1 and PP00P1_183711/1).

## Data And Code Availability

The datasets of the current study are available from the corresponding author upon request.

## Appendix

Experimental stimuli of AUT.

Experimental stimuli of AUT
|  |  |  |
| --- | --- | --- |
| Spoon | Newspaper | Sock |
| Knife | Comb | Lighter |
| Cup | Lipstick | Hat |
| Pen | Candle | Bottle |
| Toothbrush | Bucket | Hanger |
| Shoe | Clock | Tire |
| Brick | Belt | Barrel |
| Paper-clip | Newspaper | Hammer |
| Knife | Fork | Book |
| Lamp | Wine opener | Teapot |
| Spong | Basket | Trash pin |
| Screwdriver | Shop stick | Fan |
| Mirror | Tin | Umbrella |
| Scarf | Dental Floss | Toilet Paper |
| Soap | Button | Scissor |

